# A *C. elegans* Model for Spinal Muscular Atrophy Reveals Neuron Specific Developmental and Degenerative Defects

**DOI:** 10.64898/2026.01.22.701008

**Authors:** Sara Savaheli, Ivan Gallotta, Pamela Santonicola, Federica Cieri, Simon Berger, Delphine Dargere, Elia Di Schiavi, Denis Dupuy

## Abstract

Spinal muscular atrophy (SMA) is a neuromuscular disorder primarily caused by mutations in the Survival of Motor Neuron 1 (SMN1) gene. SMN1 is ubiquitously expressed and encodes a protein essential for the assembly of small nuclear ribonucleoproteins (snRNPs), key components of pre-mRNA splicing. Beyond this canonical role, SMN participates in several other fundamental cellular processes, including RNA transport, regulation of actin dynamics, transcription, and translation. While multiple hypotheses have been put forward to explain selective motor neuron (MN) vulnerability to SMN deficiency, the precise mechanisms involved remain incompletely understood. In this study, we used a simple and tractable *C. elegans* model to investigate the molecular mechanisms underlying neuronal degeneration in SMA. Silencing of the *smn-1* in targeted neurons resulted in defects in the birth and development of both motor neurons and touch receptor neurons (TRNs). In TRNs SMN-1 depletion caused distinct defects in neuronal process morphology. Our results provide evidence that key aspects of SMA pathology are conserved in *C. elegans,* which may offer new opportunities to elucidate the molecular mechanisms underlying neuronal degeneration in SMA.

## Introduction

Spinal muscular atrophy (SMA) is a genetic neuromuscular disorder first described by Guido Werdnig in 1891 ((Werdnig 1891), according to (Mercuri 2021)). SMA is characterized by the loss of motor neurons (MNs) in the anterior horn of the spinal cord and mutations of the *SMN1* gene account for over 95% of SMA cases (Crawford and Pardo 1996; Butchbach 2016). SMN protein, the product of the *SMN1* gene, is best known for its role in snRNP assembly for pre-mRNA splicing (Pellizzoni et al. 2002), but it also plays other functions, including RNA transport, cytoskeletal regulation, transcription, and translation (Singh et al. 2017). The SMN protein is ubiquitously expressed, yet its deficiency predominantly affects MNs in the spinal cord. Although SMA has more recently been recognized as a multisystem disorder involving functional impairments in multiple neuronal and non-neuronal tissue (Abati et al. 2020; Yeo and Darras 2021), neurons remain disproportionately vulnerable and represent the most prominent feature. This persistent neuronal selectivity has prompted extensive investigation into why these cells, compared to others, are uniquely affected (Simon et al. 2017; Ruggiu et al. 2012). Increasing evidence from both human and animal studies now suggests that the disorder may not arise solely from a classical neurodegenerative process. Subtle developmental abnormalities may predispose motor neurons to later degeneration (Crawford and Pardo 1996; Martínez-Hernández et al. 2013; Kong et al. 2021; McGovern et al. 2008). Additionally, the study by Irene Faravelli demonstrated dysregulated neuronal differentiation programs, supporting the need for intervention during the optimal developmental window to maximize therapeutic efficacy (Faravelli et al. 2025). Building on this emerging view, we aim to further investigate developmental alterations and their contribution to neuronal vulnerability and selective degeneration using *Caenorhabditis elegans* as an in vivo model.

*C. elegans* is widely used in biological research because it is highly amenable to genetic manipulation, possesses a simple and well-characterized nervous system, and has a fully mapped connectome, representing all neural connections (Hall and Russell 1991; White et al. 1986). These features make it an advantageous system for dissecting intrinsic, cell-autonomous neuronal differences associated with SMN loss. The *C. elegans* nervous system develops in two distinct developmental waves: of the 302 neurons that constitute the *C. elegans* hermaphrodite adult nervous system, 222 arise during embryogenesis, and the remaining are born post-embryonically during the L1 stage (Sulston et al. 1983). The GABAergic inhibitory D-type MNs form a major subgroup, further divided into six Dorsal D-type (DD) and thirteen Ventral D-type (VD) neurons. DD neurons are born during embryogenesis, while VD neurons are born at the end of the first larval stage (Sulston et al. 1983; White et al. 1986). These inhibitory MNs coordinate the alternating contraction of the dorsal and ventral longitudinal muscles, producing the animal’s characteristic sinusoidal movement. When the dorsal muscles contract, VD neurons inhibit the ventral muscles to allow relaxation; conversely, when the ventral muscles contract, DD neurons inhibit the dorsal muscles. This reciprocal pattern of activation and inhibition generates the smooth sinusoidal waves that drive forward locomotion (Howell et al. 2015; Zhen and Samuel 2015). In addition to motor control, *C. elegans* detect external stimuli using sensory neurons. Touch receptor neurons (TRNs) are a group of glutamatergic mechanosensory neurons specialized to sense gentle touch, which include the anterior lateral microtubule cells (ALMs), posterior lateral microtubule cells (PLMs), the anterior ventral microtubule cell (AVM), and the posterior ventral microtubule cell (PVM) (Allen *et al*., 2015; Schafer, 2015; Iliff and Xu, 2020). Like the D-type MNs, TRNs are generated in two distinct developmental periods: the ALM and PLM neurons arise during embryogenesis, whereas the AVM and PVM neurons are born during the first larval stage (Sulston et al. 1983).

In *C. elegans*, systemic RNAi against *smn-1* or deletion of *smn-1* results in embryonic or larval lethality and neuromuscular dysfunction but does not reliably produce overt motor neuron degeneration in surviving animals, limiting the ability to investigate the degenerative process directly (Briese et al. 2009; Miguel-Aliaga et al. 1999; Doyle et al. 2020). To overcome this, we used neuron-specific RNAi constructs (Esposito et al. 2007) to selectively silence *smn-1* in defined neuronal populations, yielding viable animals in which degeneration of targeted neurons can be studied. The RNAi constructs were driven by two distinct neuron-specific promoters—*unc-25Sp* for D-type motor neurons and *mec-3Sp* for TRNs. In this approach, RNAi is activated only in the targeted neurons and neurons develop in an *smn-1*–silenced condition (Gallotta et al. 2016). To visualize the phenotypic consequences of *smn-1* depletion, we labeled the targeted neurons by expressing fluorescent reporters driven by distinct neuron-specific promoters—*flp-13p* and *ttr-39p* for D-type motor neurons and *mec-4p* for TRNs. Neuronal morphology and development were then assessed from larval stages through adulthood.

## Results

### SMN Deficiency Alters GABAergic Motor Neuron Development

We previously determined the impact on MNs survival after *smn-1* silencing using a reporter strain in which the *unc-47* promoter drives GFP expression in all D-type MNs (Gallotta et al. 2016). In the present study, we took advantage of a reporter strain with dual labeling to reliably differentiate the effect of *smn-1* silencing on VD and DD neurons. To this end, we used the *ttr-39* promoter, driving mCherry expression in both VD and DD neurons (Jospin et al. 2009), and the *flp-13* promoter, which is active only in DD neurons to drive GFP expression (Thompson-Peer et al. 2012). As a result, VD neurons were labeled exclusively with mCherry (magenta neurons in Figure 1A–B), whereas DD neurons co-expressed both mCherry and GFP (white neurons in Figure 1A–B). Using this double reporter strain, we examined the impact of *smn-1* depletion in the two sub-classes of D-type motor neurons by reducing its expression through RNAi thanks to the strain *gbIs4* (Gallotta et al. 2016). The count of fluorescent neurons was performed from the L1 stage through the first day of adulthood and we observed a significant reduction in the total number of D-type motor neurons in the *smn-1*–silenced strain compared to the control strain (Figure 1A–B). We measured the average intensity of the 100 brightest pixels within of each individual DD neurons in 15 control and RNAi animals at the L4 larval stage (see Materials and Methods). In control animals, we observed the reporter expression to be strongest in the anterior neurons and lowest closest to the animal’s tail (Figure 1C). In SMN-1 RNAi animals we found drastic reduction of the number of detectable DD neurons, with DD1 and DD6 seeming the most sensitive to the gene perturbation. We noted that the mean fluorescence value observed for the visible neurons in the RNAi individuals was slightly higher than in the controls. This suggests that the undetectable neurons are more likely to be completely missing than present with a strongly diminished expression of the reporter. The number of DD neurons continued to decline over time (Figure 1D).

**Figure 1.**
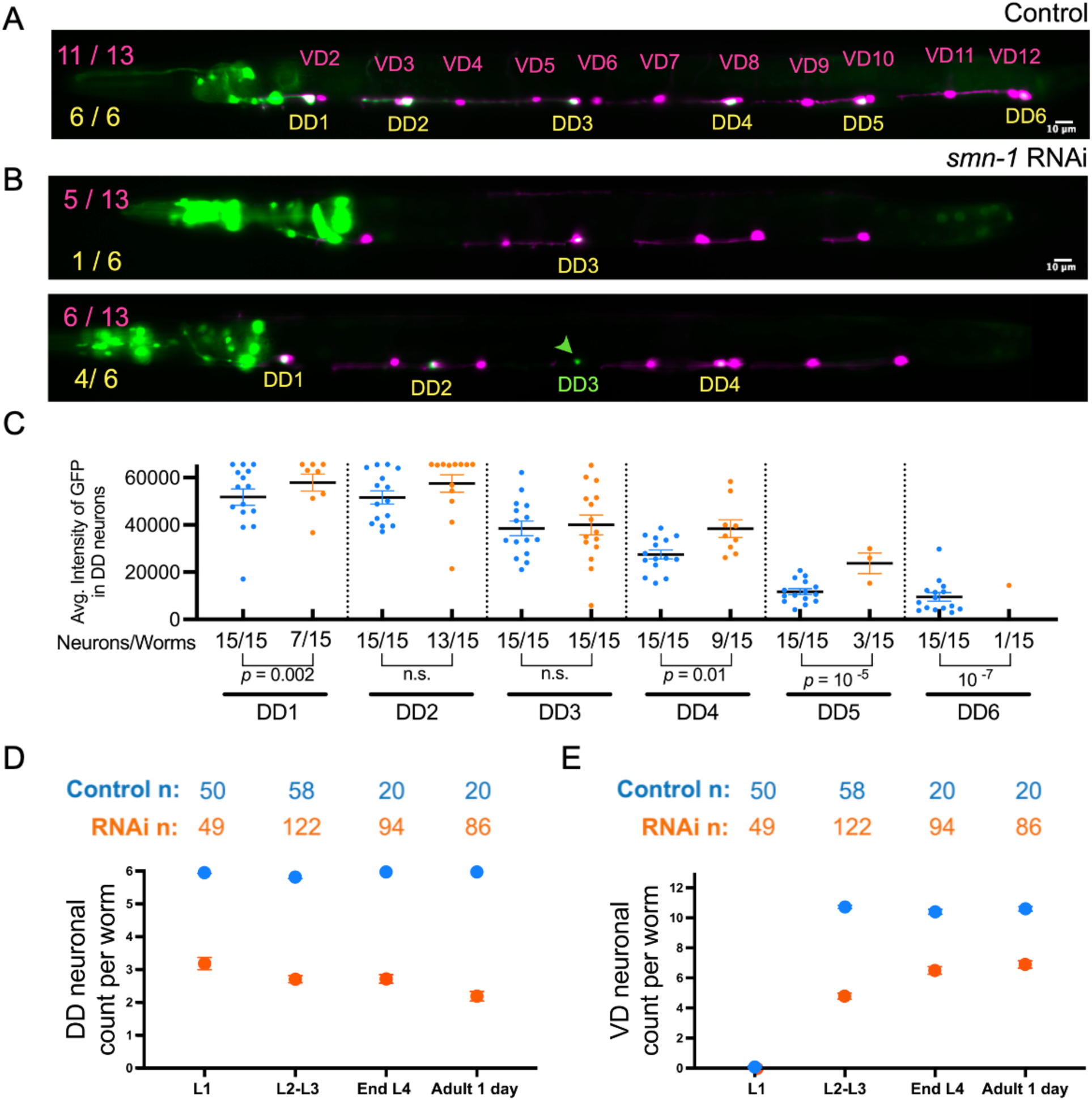
Differential effect of *smn-1* silencing on VD and DD Neurons. (A) Representative image of VD and DD motor neurons in the control reporter strain (DUD2204) at the end of the L4 larval stage carrying *flp-13p*::GFP (expressed in DD neurons) and *ttr-39p*::mCherry (expressed in VD and DD neurons). Eleven out of thirteen VD neurons (magenta) and all six DD neurons (white) are visible. (B) Representative images of two animals (strain DUD2205) at the end of the L4 larval stage, carrying *unc-25Sp::smn-1 RNAi (smn-1* silencing), *flp-13p*::GFP (GFP expressed in DD neurons) and *ttr-39p*::*mCherry* (mCherry expressed in VD and DD neurons). Upper image: five out of thirteen VD neurons (magenta) and only one DD neuron (white) are visible. Lower image: six out of thirteen VD neurons (magenta) and four out of six DD neurons (white) are visible. One of the DD neurons (green arrowhead) expresses only GFP instead of the expected co-expression of GFP and mCherry. (C) Comparative analysis of *flp-13p*::GFP reporter intensity in each DD neuron at L4 larval stage in the control (blue dots) and in *smn-1*–silenced strains (orange dots). Below the x-axis are noted the ratio of detectable neurons on the total number of worms analyzed. Significance of neuronal counts difference between conditions were evaluated using Fisher’s exact test, when significant, exact *p* values are indicated below the brackets. (D and E) Comparative analysis of motor neurons in control and *smn-1*–silenced strains by counting visible DD neurons (D) and VD neurons (E) in the control (blue dots) and in *smn-1*–silenced strains (orange dots) at different time points. Head neurons fluorescence correspond to the co-injection markers *ttx-3p::GFP* and *chs-2p::GFP,* indicating the presence of *ttr-39p*::*mCherry* and the *smn-1* silencing construct respectively.

The VD neurons appear during the L2 stage, they were thus absent at L1 stage in both control and *smn-1*–silenced strains, both control and *smn-1*–silenced strains (Figure 1E). By the L2 stage, when the full complement of VD neurons had emerged in control worms, *smn-1*–silenced animals displayed a marked reduction that persisted through adulthood, indicating a developmental deficit. We observed a slight increase in neuronal counts during post embryonic development (Figure 1E).

We used long-term time-lapse imaging (Berger et al. 2021) to monitor the birth of VD neurons during the transition from L1 to L2 in *smn-1*–silenced animals and confirmed that some VD neurons failed to appear (see supplementary data, Video 1-2). In rare cases, the mCherry signal was absent in some DD neurons, (Figure 1B, arrowhead; Supplementary Video 1) suggesting that *ttr-39*-promoter–driven mCherry expression might be affected by reductions in SMN-1 levels. This observation reinforces the need for caution when interpreting the absence of a fluorescent signal: the lack of visible expression may indicate the failure of the targeted cell to express its fluorescent marker rather than the absence of the expected neuron.

We measured the average intensity of the 100 brightest pixels within each neuron in mCherry fluorescent channel at the L4 larval stage (see Materials and Methods). We did not observe a pronounced change in *ttr-39p*::mCherry reporter levels in either DD or VD neurons in *smn-1*– silenced animals compared to controls (see supplementary Figure S1).

These findings suggest that the *smn-1* silencing produced developmental defects in VD neurons and the absence of mCherry in a subset of DD neurons likely reflects major transcriptional changes in the targeted neurons.

### SMN Deficiency Alters Touch Receptor Neurons Development and Survival

To determine whether the developmental defects observed in VD neurons are specific to those neuron types, we tested SMN-1 depletion in Touch Receptor Neurons (TRNs). For visualization of the TRNs, we used a GFP label, driven by the *mec-4* promoter that also enables clear visualization of their structure and morphology (Figure 2A). We crossed the TRN reporter strain with strain *dudIs2202* in which *smn-1* is selectively silenced in TRNs (Figure 2B). Imaging was performed from the L1 larval stage through young adulthood. Quantification of ALM and PLM neurons in the *smn-1*–silenced strain revealed that, although these embryonically derived neurons were present in normal numbers at the L1 stage, the GFP reporter intensity was markedly lower than control. As development progressed, GFP reporter intensity declined even further, associated with a reduced number of detectable neurons (Figure 2C-D).

**Figure 2.**
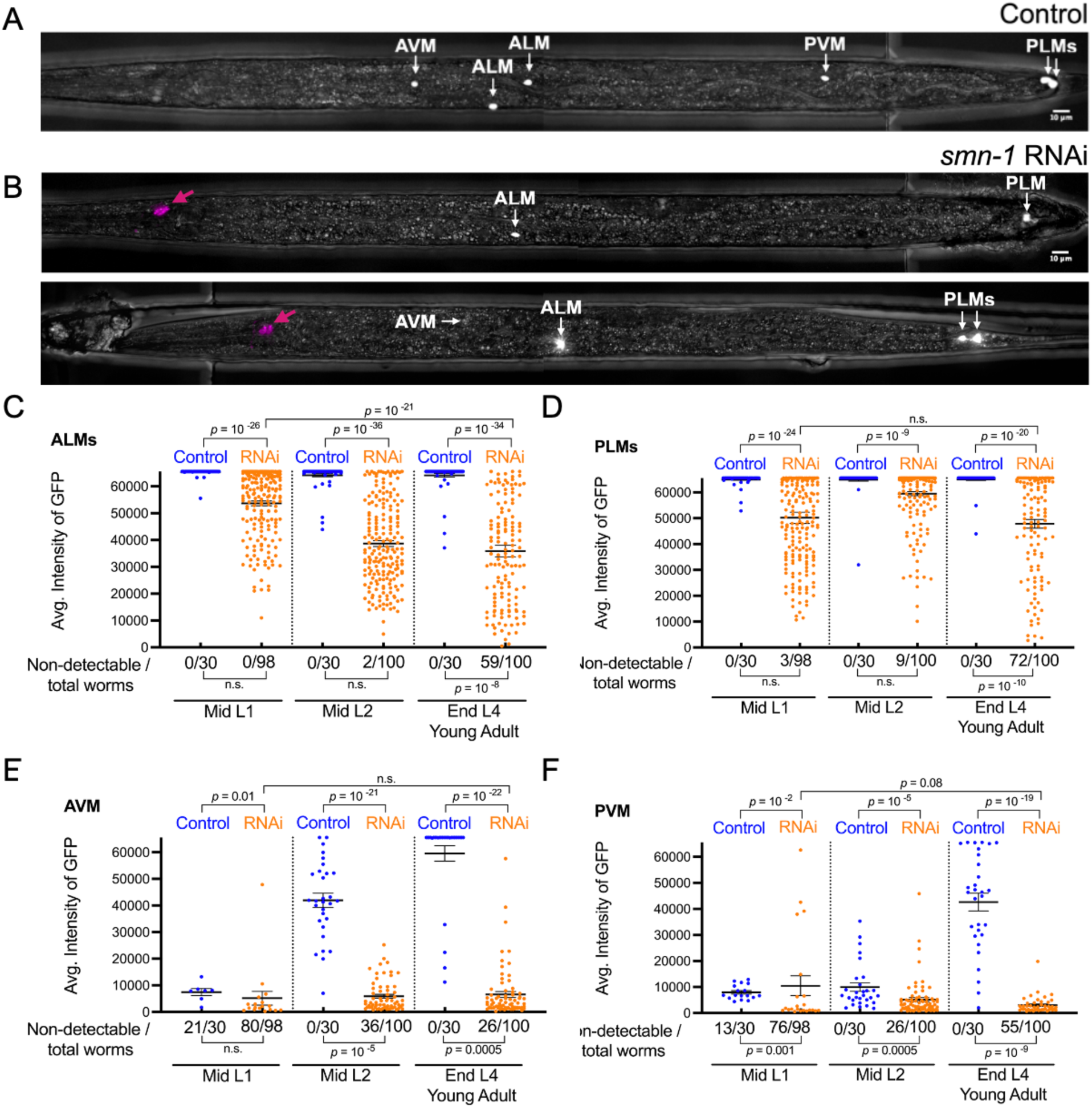
Distinct effects of *smn-1* silencing on TRNs neuronal development and maintenance. (A) Representative images of TRNs in the control strain carrying *mec-4p*::GFP reporter at the end of the L4 larval stage. All six TRNs are visible, as indicated by the white arrows. (B) Representative images of two *smn-1*–silenced animals (strain NA2397) carrying *mec-4p::*GFP and *mec-3Sp::smn-1 RNAi* transgenes, at the end of the L4 larval stage. Visible TRNs are indicated by white arrows. The number of visible TRNs is reduced to 2 in the upper image and 4 in the lower image. The magenta arrow indicates the co-injection marker. (C–F) Quantification of neuronal intensity (control: blue dots; *smn-1*–silenced: orange dots) at three different developmental timepoints (mid L1, mid L2 and end of L4/young adulthood). The frequency of non-detectable neurons is noted below the x-axis as the number of non-detectable neurons over the total number of worms analyzed. Differences in neuronal intensity and neuronal counts between conditions were statistically evaluated. When significant, exact *p* values are displayed above or below the bracket.

We next examined the developmental progression of AVM and PVM neurons. In the control strain, these neurons were not detectable in early L1 as expected but became visible in all worms by the L2 stage. GFP reporter signal intensity increased progressively during development, reaching maximal levels by adulthood (intensities: 59,557 ± 2,879 for AVM and 42,637± 3,410 for PVM). In contrast, in the *smn-1*–silenced strain, a subset of animals failed to develop detectable AVM and PVM neurons by the L4–young adult stage. In neurons that were present, the reporter intensity remained markedly reduced (6,584 ± 1,101 for AVM and 3,018 ± 473 for PVM) (Figure 2E–F). Additionally, we observed a reduction of the number of visible PVM neurons which were not visible in 26 out of one hundred mid-L2 RNAi animals. This number rose to 55 at the L4–young adult stage (Figure 2F). In contrast, AVM neurons did not show this phenotype (Figure 2E) indicating differential sensitivity between the two neuron types. Overall, our observations indicate that *smn-1* silencing affects both the development and survival of TRNs.

### Structural Abnormalities in ALM and PLM Neuronal Processes Under *smn-1* Silencing

ALM and PLM provide an ideal system for studying morphological defects in the neurite process. ALM neurons are positioned in the midbody and extend a long anterior process that branches ventrally into the nerve ring (see supplementary Figure S1). In a subset of animals, ALM neurons also exhibit a short posterior protrusion extending less than twice the length of the cell body. PLM neurons on the other hand are located in the lumbar ganglion, extending a long anterior process that terminates near the ALM cell body, and a short posterior process (Chalfie et al. 1985; Kirszenblat et al. 2013). We quantified the size of the anterior process in ALM and PLM neurons after hatching and at subsequent larval stages in control and *smn-1 RNAi* animals. The neurite processes were severely shortened in nearly all *smn-1*–silenced animals (Figure 3B). At the L1 stage, ALM and PLM anterior processes in *smn-1*–silenced worms are markedly shorter than those in control worms (Figure 3C, D). However, nearly all ALM and PLM neurons were still present in the *smn-1*–silenced worms at the L1 stage (Figure 2C-D). Throughout development, while the ALM neurite process in control animals grows steadily, in *smn-1*– silenced worms they remained markedly shorter despite a slight increase in length (Figure 3B-D). We also detected ALM neurons lacking a neurite process at the L4 stage in *smn-1*–silenced worms, indicating not only a failure to extend the ALM neurite but also that some defective neurites show retrograde degeneration in an age-dependent manner (Figure 3C). In the PLM neurons instead there was no measurable growth of the neurite over time (Figure 3C, D), once again uncovering distinct phenotypes between different neurons.

**Figure 3.**
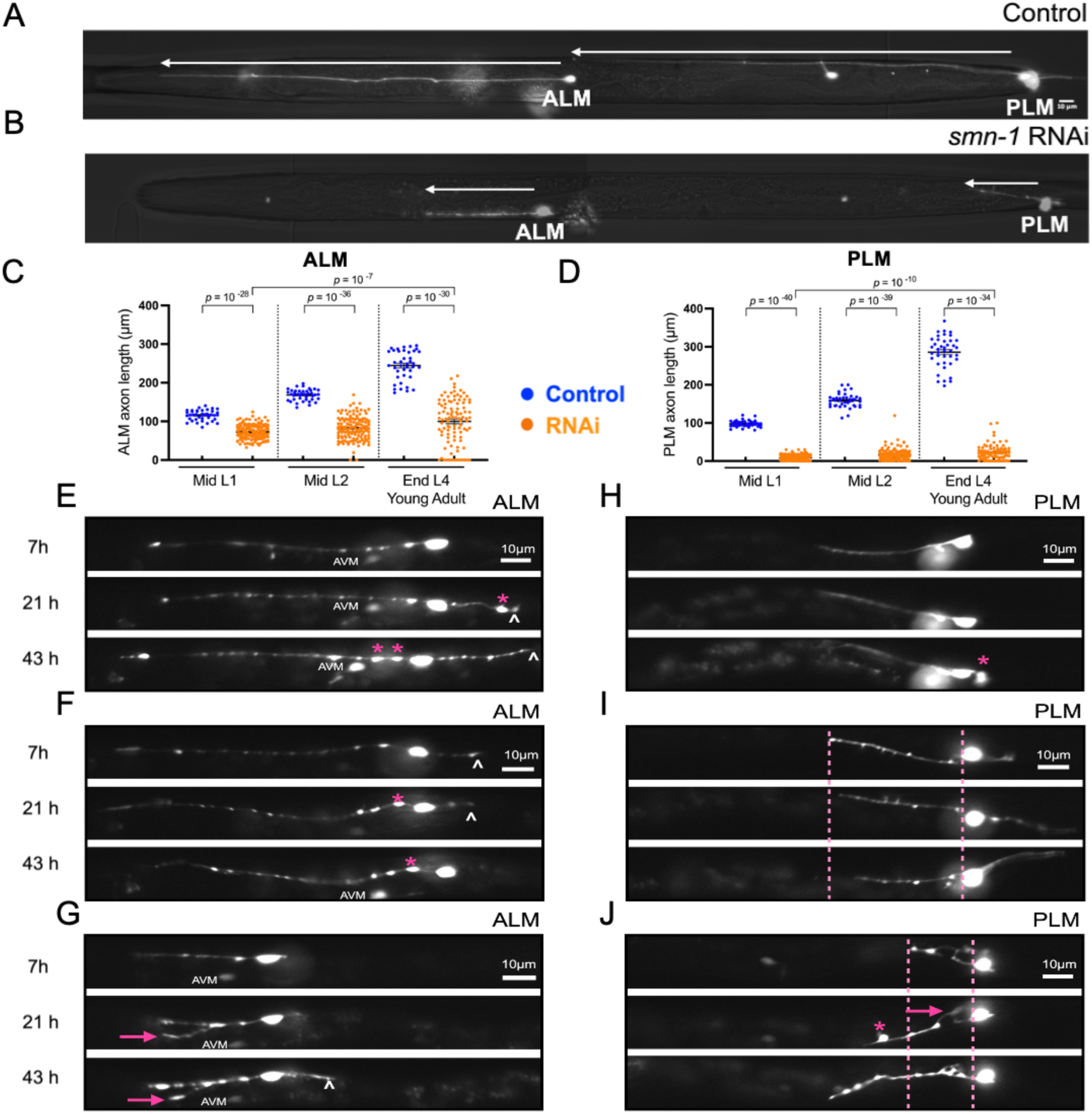
Degeneration of ALM and PLM neurites in *smn-1*–silenced *strain*. **(A)** Representative image of the neuronal processes of ALMs and PLMs in the control strain carrying *mec-4p::*GFP at the end of the L4 larval stage. (B) Representative image of neuronal processes of ALMs and PLMs in the *smn-1*–silenced strain carrying *mec-4p::*GFP and *mec-3Sp::smn-1 RNAi* at the end of the L4 larval stage. Quantification of ALM and PLM neurite lengths (μm) was performed across different developmental stages. (C, D) Comparative analysis of ALM (C) and PLM (D) neuronal process length in control (blue dots) and *smn-1*–silenced strains (orange dots). A total of 20 control worms and 70 *smn-1*–silenced worms were examined in this study. Significance of neurite length distributions difference between conditions were statistically evaluated. When significant, exact *p* values are displayed above the horizontal bracket. (E–J) Representative image of neuronal process abnormalities in *smn-1*– silenced strain. Ectopic posterior neurite elongation in ALM neuron (white arrowhead) (E-G). Bulging in the neuronal process (magenta star) of ALM (E-G) and PLM (H, J). Abnormally branched neurites in ALM (G) and PLM (J). Progressive shortening of the anterior neurite in PLM (I) and abnormal dynamic of the anterior neurite in PLM (J).

Through long-term time-lapse imaging experiments (Berger et al. 2021), we consistently observed short anterior processes across all examined neurons, whereas neurite branching was detected only in a subset of these shortened processes. In neurons that developed branches, these structures were initially absent and emerged during development (Figure 3G, ALM; Figure 3J, PLM). Additionally, we observed elongated ectopic posterior neurites in certain ALM neurons that progressively extended over time (Figure 3E, G), whereas in others, existing ectopic neurites gradually shortened and eventually disappeared (Figure 3F).

Another notable pattern is the presence of focal swellings and bulges along the length of the neuronal processes under *smn-1* silencing condition (Figures 3E–H, J). These swellings mildly increased in size over time. Processes affected by such swellings were eventually lost, followed by disappearance of the corresponding neuronal cell body, indicating that neurite disappearance precedes neuronal loss (see supplementary data, Video 3). Overall, our findings showed the presence of neuronal process defects preceding neuronal degeneration.

## Discussion

The SMN is a multifunctional protein with essential roles in RNA processing and additional roles in diverse cellular processes (Singh et al. 2017). SMN is broadly expressed across tissues, and while severe SMA can affect multiple cell types, including sensory neurons, MNs appear to be more vulnerable to SMN depletion (Anagnostou et al. 2005; Jablonka et al. 2006; Abati et al. 2020; Yeo and Darras 2021). Several studies have investigated the mechanisms underlying this selective vulnerability, since understanding these mechanisms could guide the development of complementary and more targeted therapeutic strategies (Ruggiu et al. 2012; Simon et al. 2017). Extending these efforts, in this study we employed a *C. elegans* model of SMA to track SMA-associated neuronal alterations on D-type MNs and TRNs.

Earlier studies have shown that *smn-1* silencing in *C. elegans* GABAergic MNs, including VD and DD subtypes, significantly compromise the survival of neurons in an age-dependent manner (Gallotta et al. 2016). Here, we show that *smn-1* silencing also affects development of the studied VD neurons, with defects observed in the appearance of a subset of these neurons (Figure 1E). To determine whether this phenotype was specific to MNs, we examined TRN mechanosensory neurons. We observed comparable defects in TRNs, which was not unexpected, given that sensory neurons also become compromised in more severe cases in human and animal studies (Anagnostou et al. 2005; Jablonka et al. 2006). These findings indicate that sensory neurons are not immune to the consequences of SMN-1 loss and may exhibit dysfunction as part of the disease cascade. Analysis of the observed phenotype revealed that *smn-1* silencing via RNAi caused severe defects in AVM and PVM neuron development, with these neurons either absent or showing markedly reduced *mec-4p*::GFP reporter expression (Figure 2E, F). Considering that *mec-4* gene *is* a terminal differentiation marker (Zheng et al. 2018), strong reduction of *mec-4p*::GFP expression could indicate that *smn-1* depletion is severely impairing proper AVM and PVM differentiation.

Developmental defects in *C. elegans* VD, AVM and PVM neurons – are reminiscent of previous findings from human and animal studies, indicating that SMA is better understood as a developmental defect, rather than purely a degenerative condition, where impaired neuronal development predisposes cells to future degeneration (Crawford and Pardo 1996). Additionally, studies in zebrafish, fruit fly, and mouse models have reported SMN-1 depletion causing defects to neurogenesis, axonal development, and neuromuscular junction (NMJ) maturation, indicating that there are multiple effects converging on the pathology (Martínez-Hernández et al. 2013; Kong et al. 2021; McGovern et al. 2008). In our study, the developmental defects observed in the studied neurons suggest that *smn-1* silencing disrupts their normal maturation and prevents these neurons from achieving their full differentiation state, indicating an impairment in neurogenesis. Beyond these developmental abnormalities, we observed an age-dependent loss of TRNs, predominantly affecting the embryonically derived ALM and PLM neurons (Figure 2C, D). These neurons progressively lost *mec-4p*::GFP expression in *smn-1*-silenced animals, which may reflect a failure to maintain terminal identity or premature neuronal degeneration. Consistent with this interpretation, age-dependent neuronal loss was observed in post-embryonically born PVM neurons that had exhibited earlier developmental defects (Figure 2F). In contrast, post-embryonic VD and AVM neurons did not display a comparable age-dependent degenerative phenotype (Figure 1E, 2E).

We also detected progressive structural changes in ALM and PLM neuronal processes, such as shortening and the emergence of branched neurites (Figure 3E-J). Another notable phenotype observed is ectopic enlargement of the posterior neurites in some ALM neurons (Figure 3E-G), a similar phenotype observed in mutants for *mec-7* (*ky852)* (Kirszenblat et al. 2013) and *ptrn-1* (Chuang et al. 2014) genes, both being essential for microtubule dynamics. Evidence for defective microtubule stabilization and polymerization has been reported in SMA models and is attributed to the upregulation of Stathmin, a known microtubule-destabilizing factor (Wen et al. 2010). In general, neuronal process branching and ALM posterior neurite appearance observed in our model therefore suggest a defect of microtubule integrity in TRNs, leading to neuronal process abnormalities reminiscent of those observed in other microtubule-defective mutants.

We also identified the presence of neurite swelling all along the neurite of some TRNs when SMN-1 is depleted (Figures 3E–H, J). These focal axonal swellings increase in size and number over time. This type of axonal swelling has previously been reported in both human and animal SMA models ((Coer̈s and Woolf 1959) according to (McGovern et al. 2008)) and other neurodegenerative conditions, such as spastic paraplegia and has been attributed to impaired axonal transport (Tarrade et al. 2006). Given that one of the SMN protein’s known functions is in axonal transport (Singh et al. 2017), it is plausible that disruptions in axonal transport or even microtubule integrity might contribute to the appearance of these swellings. However, further studies are needed to explore the exact mechanisms behind these neurite swellings and their role in SMA progression.

Finally, we observed changes in the expression of the neuron-specific reporters used in this study. In TRN neurons, *smn-1* silencing resulted in a generalized reduction in GFP expression driven by the *mec-4* promoter (Figure 2C–F), suggesting altered regulation or maintenance of *mec-4* gene expression. In motor neurons, we occasionally detected a loss of mCherry fluorescence driven by the *ttr-39* promoter in some DD neurons, despite robust GFP expression from the *flp-13* promoter that confirms neuronal presence (Figure 1B, Green arrowhead). Notably, GFP expression driven by the *flp-13* promoter was mildly increased in *smn-1*-silenced animals compared with controls (Figure 1C), which warrants further investigation to determine if loss of SMN-1 influences *flp-13* transcript abundance and protein levels. Since SMN’s primary role is in RNA metabolism, *smn-1* silencing is expected to cause widespread transcriptional disruption. The observed defect in reporter expression may reflect either neuronal death or an altered transcriptional profile that disrupts terminal differentiation, or neuronal identity (Corti et al. 2012; Lotti et al. 2012; Bäumer et al. 2009).

This study also highlighted several instances of differential phenotypes between neurons of the same class: from distinct penetrance and intensity of reporter expression between different MNs to different phenotypic responses to RNAi between ALM and PVM. This later observation could either be the consequence of a different level of activation of the *mec-3* promoter driving the RNAi in these cells or a differential sensitivity to the SMN-1 depletion in these cells.

In conclusion, our findings revealed defects in both neuronal development and survival, accompanied by specific neurite abnormalities, which collectively mimic the phenotypes of SMA observed in established SMA models. The defects observed in our model further underscore the utility of *C. elegans*, for detailed investigation of the molecular mechanisms underlying the SMA phenotype.

## Material And Methods

### *C. elegans* Strains and Culture Conditions

Worms were grown at 20°C onto NGM plates seeded with bacteria (*Escherichia coli*, OP50) as previously described (Brenner 1974). In order to maintain the population, worms were transferred onto new plates every 3 days. Wild-type animals were *C. elegans* variety Bristol, strain N2. The alleles and transgenic strains used in this work are summarized in Supplementary Table S1.

### Worm synchronization

Worms were cultured for 2–3 days on seeded NGM plates to obtain gravid adults. Gravid adults were bleached in a solution of sodium hypochlorite (commercial bleach) and NaOH (final concentrations: NaOH 1 M, sodium hypochlorite 0.4–1.6%) and vortexed for 5 minutes. Once most of the worms had dissolved, bleaching was stopped by adding M9 buffer. The eggs were then pelleted and washed three times with M9 buffer to remove any residual bleach. Finally, the eggs were distributed onto seeded NGM plates and incubated at 20 °C to allow further development.

### Neuron-specific RNAi knock-down

To knock-down *smn-1* specifically in TRNs we used a short form of *mec-3* promoter (417 bp) (Toms et al. 2001) to drive the exon-rich region used for the RNAi feeding library (Kamath et al. 2003). Exon-rich regions were amplified in two separate PCR reactions to obtain the sense and antisense fragments that were fused to the *mec-3* promoter by PCR fusion using internal primers, as previously described (Esposito et al. 2007; Hobert 2002).

### Transgenic lines

Germline transformation was performed as described previously (Mello et al. 1991). Sense and antisense fusions to the TRNs promoter were pooled in equimolar amounts and microinjected in the N2 strain at 50 ng/μL concentration, to obtain *gbEx518* transgene: *GBF308 mec-3Sp::smn-1 (RNAi sense/antisense); odr-1p::RFP*. The co-injection marker used in this work was: *odr-1p::RFP* (kindly provided by C. Bargmann, The Rockfeller University, New York, USA; RFP expression in AWC and AWB neurons). The injection marker was co-injected at 30 ng/µl.

### Integration of Extrachromosomal Arrays by ultraviolet irradiation

The injected DNA molecules rearrange to form multicopy extrachromosomal arrays, which can be unstable or mosaic. Integration of these extrachromosomal arrays into the genome provides a means to achieve stable and uniform expression in the progeny, typically through irradiation with ultraviolet or gamma rays (Mariol 2013). In this study, we used ultraviolet irradiation to integrate one of the extrachromosomal array, *gbEx518a* transgene, carried by strain NA1164 to generate the stable integrated transgenic strain DUD2202, with the genotype *dudIs2202* [*mec-3Sp::smn-1 (RNAi sense/antisense*); *odr-1p::RFP*] III. To identify the chromosome into which the extrachromosomal array was integrated, we employed *C. elegans* mapping strains.

The integration procedure was performed following the protocol described by Mariol *et al*. (Mariol 2013) with minor modifications. Worms carrying an extrachromosomal array with an estimated transmission rate of 20–30% were used. Fifteen L4 or young adult hermaphrodites were transferred onto 10 cm NGM plates seeded with *E. coli* OP50. The plates were irradiated using a UV crosslinker (UV 701435, Jeulin) equipped with a 254 nm UV lamp positioned 5 cm above the plate surface. Worms were exposed to UV light at an intensity of 0.012 J/cm² for 100 seconds with the plate lids removed. Following irradiation, plates were incubated at 20 °C to allow the worms to recover and lay eggs. When the progeny reached the young adult stage, fluorescent co-injection markers were used to identify transgenic individuals. Ninety fluorescent F₁ animals were isolated and transferred individually onto separate 60 mm NGM plates seeded with *E. coli* OP50. These single F₁ worms were incubated at 20 °C until their progeny reached the young adult stage to allow for observation of fluorescent F₂ animals. Plates showing no progeny or a transmission rate below 30% were discarded. From each selected F₁ plate, four highly fluorescent transgenic F₂ animals were isolated and transferred individually onto new 60 mm NGM plates. The resulting F₂ progeny were subsequently screened for 100% fluorescent F₃ worms, indicative of successful integration of the transgene.

### Genetic crosses

Hermaphrodite and male worms were transferred to a fresh NGM plate at a ratio of 5:2, along with a drop of *E. coli* OP50 to promote proximity and facilitate mating. A successful cross was identified by the presence of the paternal fluorescent marker in the F₁ progeny. Four F₁ worms exhibiting both paternal and maternal fluorescent markers were isolated and allowed to produce F₂ progeny. From each F₁ plate, eight F₂ worms were transferred individually onto separate NGM plates, and their progeny were examined for the presence of 100% expression of both paternal and maternal fluorescent markers, confirming successful establishment of double transgenic lines.

### Microscopy Observations

Before starting the observations, a 2% agarose pad was prepared on a glass microscope slide. A drop of 10 mmol/L levamisole or sodium azide was applied to the pad to immobilize the worms without causing lethality. Worms were then transferred into the anesthetic solution, and a coverslip was gently placed on top. Observations were performed using a Zeiss Axio Imager Z1 microscope equipped with an HXP 120 lamp and an Axiocam MRM camera for image acquisition. Worms were examined using 10× or 40× objectives under DIC (Differential Interference Contrast) optics, followed by fluorescence imaging (excitation at 395 nm for GFP and 558 nm for dsRed). Neurons were counted manually, and each transgenic strain was analyzed independently rather than through direct comparison of phenotypic outcomes between the two neuronal populations, because the promoters used to drive RNAi expression in D-type motor neurons (*unc-25p* short) and TRNs (*mec-3p* short) differ, and the extent of *smn-1* silencing may therefore vary between neuronal subtypes.

Long-term imaging experiments were performed as described in Berger et al. 2021 and 2025 (Berger et al. 2021, 2025). Briefly animals were loaded into the stage appropriate microfluidic long-term imaging device and continuously fed with E. coli NA22. Animals were imaged at 15 min intervals for up to 48 hours. All images were acquired using an epifluorescence microscope, consisting of a sCMOS camera, an LED light source for fluorescence and brightfield illumination and a piezo objective drive for Z-motorization, using a 40×/1.25NA oil immersion lens. Image acquisition and actuation of the on-chip hydraulic valve was coordinated using a custom MATLAB script. All images were acquired at 20±0.5°C, with temperature control achieved either via the room air-conditioning system, or a microscope cage incubator. Large numbers of animals were imaged as described in Spiri et al. 2022 (Spiri et al. 2022), with animals grown on NGM plates until the desired stage is reached. At this point animals are washed off the NGM plate and loaded into a stage appropriate imaging device and anesthetized using levamisole. Animals were then imaged using an epifluorescence microscope, likewise equipped with a sCMOS camera, an LED light source for fluorescence and brightfield illumination and a piezo objective drive for Z-motorization. Images were acquired using a 40×/1.1NA water immersion lens, and acquisition was coordinated using a custom MATLAB script.

### Quantification of Fluorescence Intensity

For quantification of fluorescence intensity, we measured the average intensity of the 100 brightest pixels within the D-type motor neuron and TRN neuron. The pixel intensity ranges from 0 to 65,535 in 16-bit images.

For the D-type motor neurons, we first generated composite images for each worm in FIJI to distinguish VD and DD neurons. This produced three composite images per worm, with each composite correctly representing the intensity of one reporter or the bright-field image. To quantify fluorescent intensity for each reporter in VD or DD neurons, measurements were performed on composite images in which the corresponding reporter channel was set as the primary channel. For TRNs, the creation of composite images was omitted, and intensity measurements were taken directly from the TRN reporter channel. Fluorescence intensity measurements were performed using *Fiji/ImageJ* (version 2.16.0/1.54p). For D-type motor neurons and TRNs, two different codes were used to measure the average intensity of the 100 brightest pixels due to differences in the anatomical positions of the neurons in the worms. For both neurons, a region of interest (ROI) was first manually defined around each neuron in the visually optimal optical slice.

For D-Type Motor Neurons, it is not expected that more than one neuron would appear within the defined ROI across different optical slices because D-Type Motor Neurons are anatomically positioned adjacent to one another in the horizontal plane. Therefore, the analysis was extended across all optical slices to identify the slice exhibiting the maximum signal. Using a macro, pixel intensity values within each ROI were extracted across all 41 slices using the **getRawStatistics** function. Then, the average intensity of the 100 brightest pixels was calculated for each slice. The maximum of these average values across all slices was then reported for each ROI in the Results table and used for subsequent analyses.

For TRNs, an alternative code with the same purpose was used, since it is possible for two neurons to be present within the same ROI in different optical slices. In this code, after manually defining an ROI around each neuron in the visually optimal optical slice, the script automatically searched within a range of ±5 optical slices around the selected focal plane to identify the best slice exhibiting the highest mean fluorescence intensity within the ROI. In this best slice, pixel intensity values within the ROI were extracted using the **getRawStatistics** function. The script then sorted all pixel values in descending order and calculated the average intensity of the 100 brightest pixels for each neuron (or fewer when fewer than 100 pixels were available). The resulting average value was recorded for each analyzed ROI and reported in the Results table for subsequent analysis.

The full *Fiji* macro code used for these quantifications is provided in the Supplementary Materials.

### Statistical analysis

Statistical analyses were performed using GraphPad Prism 10 and Python (v3.10.19; SciPy and pandas). In all experiments, independent worms were analyzed at each time point. Data normality was assessed using the Shapiro–Wilk test, and datasets with *p* ≥ 0.05 were considered normally distributed. Neuronal intensity distributions and neurite length distributions were compared pairwise between conditions using the non-parametric Mann–Whitney U test. Neuronal count data were analyzed pairwise between conditions using Fisher’s exact test.

## Acknowledgements

The *CZ8332 and SK4005* nematode strains used in this work were provided by the *Caenorhabditis* Genetics Center, CGC (funded by NIH Office of Research Infrastructure Programs P40 OD010440). The authors thank C. Bargmann (The Rockfeller University, New York, USA) for *odr-1p::RFP* containing plasmid, Y. Jin and A. Chisholm for CZ2475 strain, Wormbase (Version: WS298), BioRender, and would like to acknowledge Giuseppina Zampi, Antonio Suppa and Marco Petruzziello (IBBR, Napoli) for technical support, and Chiara Nobile (IBBR, Napoli) for administrative support.

## Fundings

This work was supported by: Vinci program (2021); the Italian Telethon Foundation (project number #GGP16203) to EDS; the AFM-Telethon (project number #24401) to EDS; the National Recovery and Resilience Plan (NRRP), Mission M4C2I1.3 (funded by the European Union – NextGenerationEU, Project code PRR.AP015.090, CUP E63C22002170007) to EDS.

## Author contributions

Study concepts and design: EDS, DDu; *C. elegans* experiments and data evaluation: SS, PS, IG, FC, DDa, SB; manuscript preparation: SS, EDS, DDu; approval of the final version of the manuscript submitted: all authors.

## Ethics declarations

The authors declare no competing interests.

## Supplementary Materials

### *FIJI* macro code for D-Type Motor Neurons and TRNs

For D-Type Motor Neurons, a macro code was used to calculate the average intensity of the 100 brightest pixels within the defined ROI across all optical slices, and only the maximum average value per ROI was reported and was used for subsequent analysis.

For TRNs, a different macro code was used due to their positional characteristics, as searching across all optical slices could result in the inclusion of signals from neighboring neurons within the defined ROI. Therefore, this code first identified the slice with the highest mean intensity within the ROI across a range of ±5 slices around the focal plane and then calculated the average intensity of the 100 brightest pixels in that slice. This value was reported and used for subsequent analysis (for a detailed description, see the Materials and Methods section).

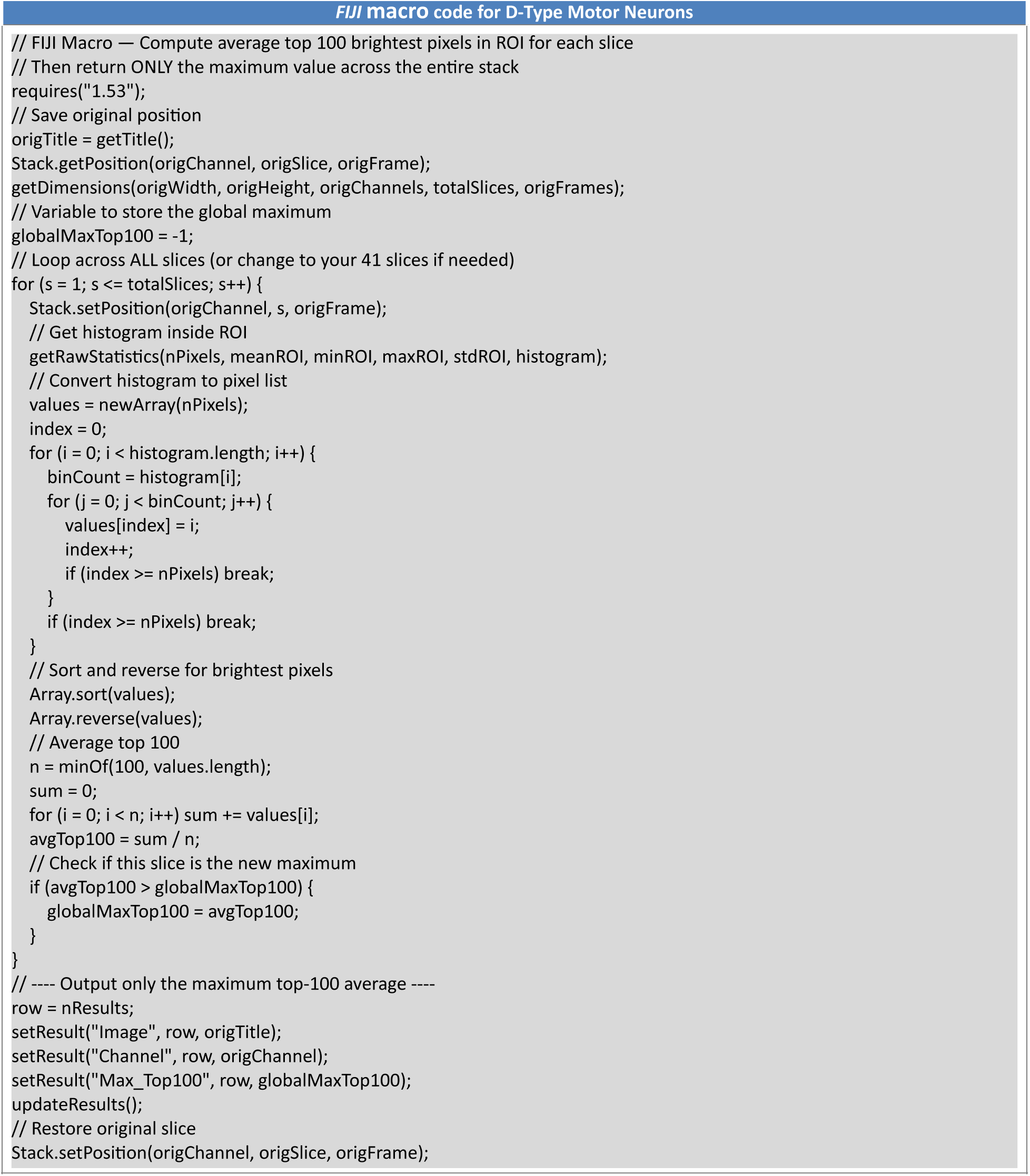

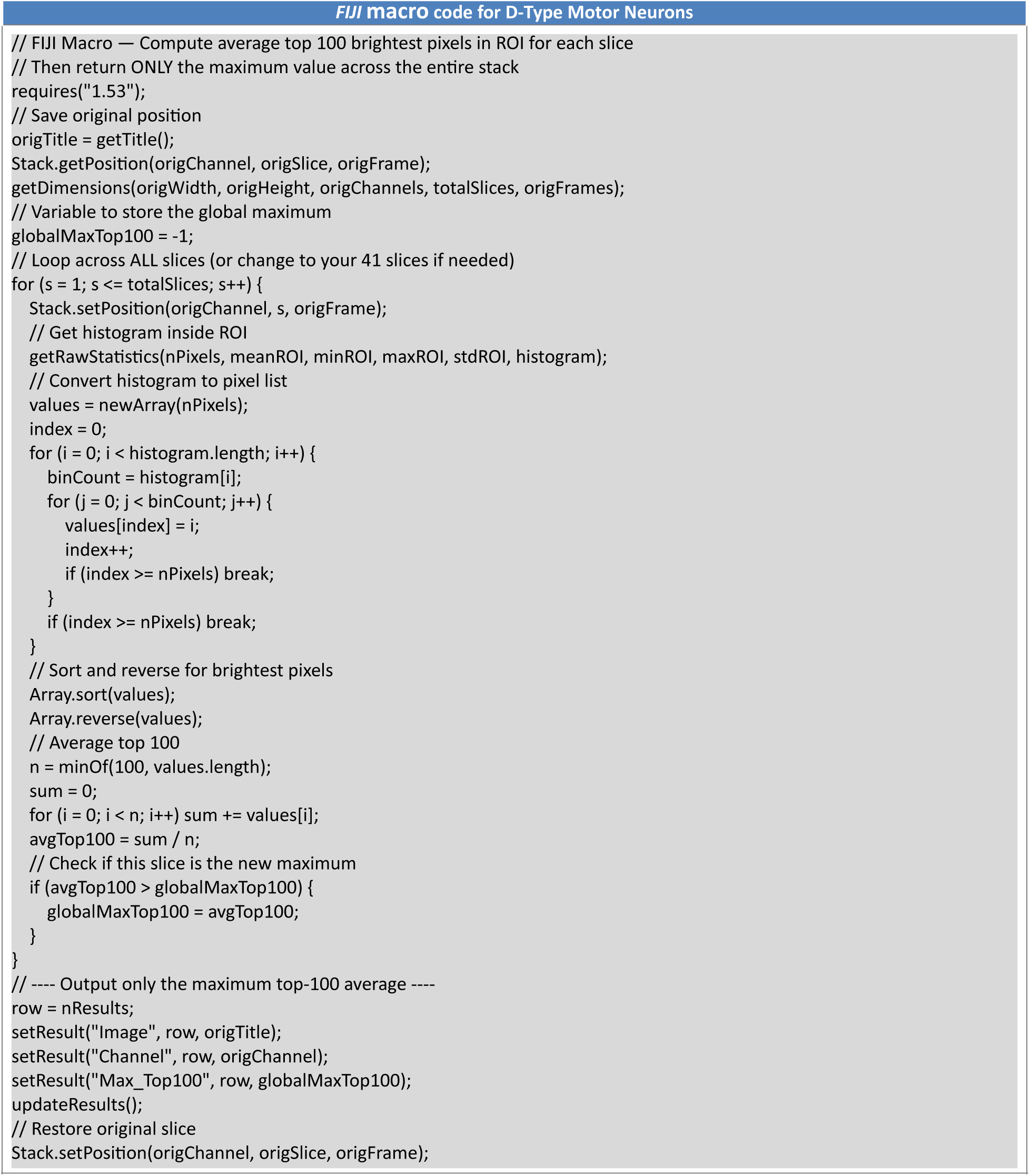

**Table S1.**
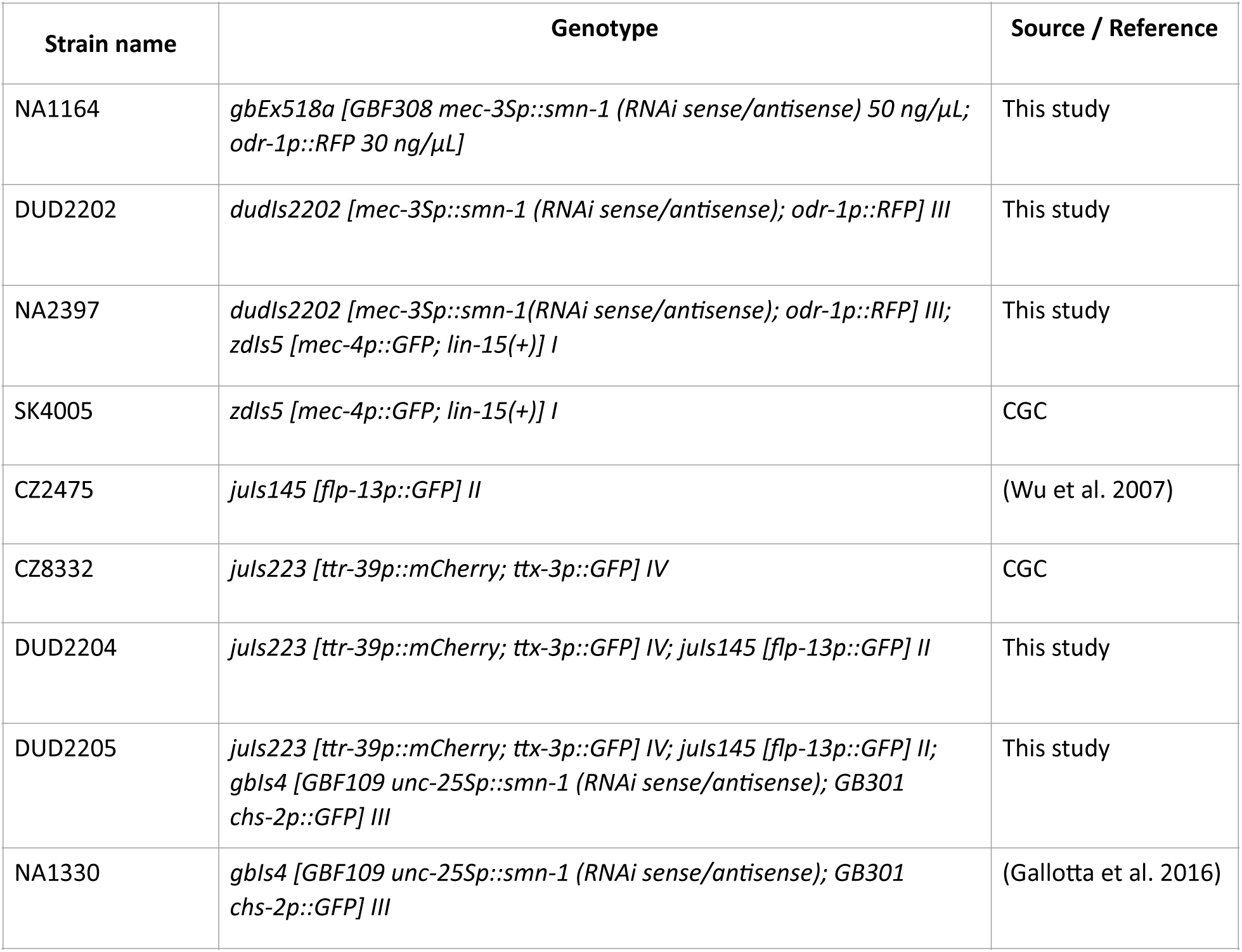
Strains used in this study.

**Figure S1.**
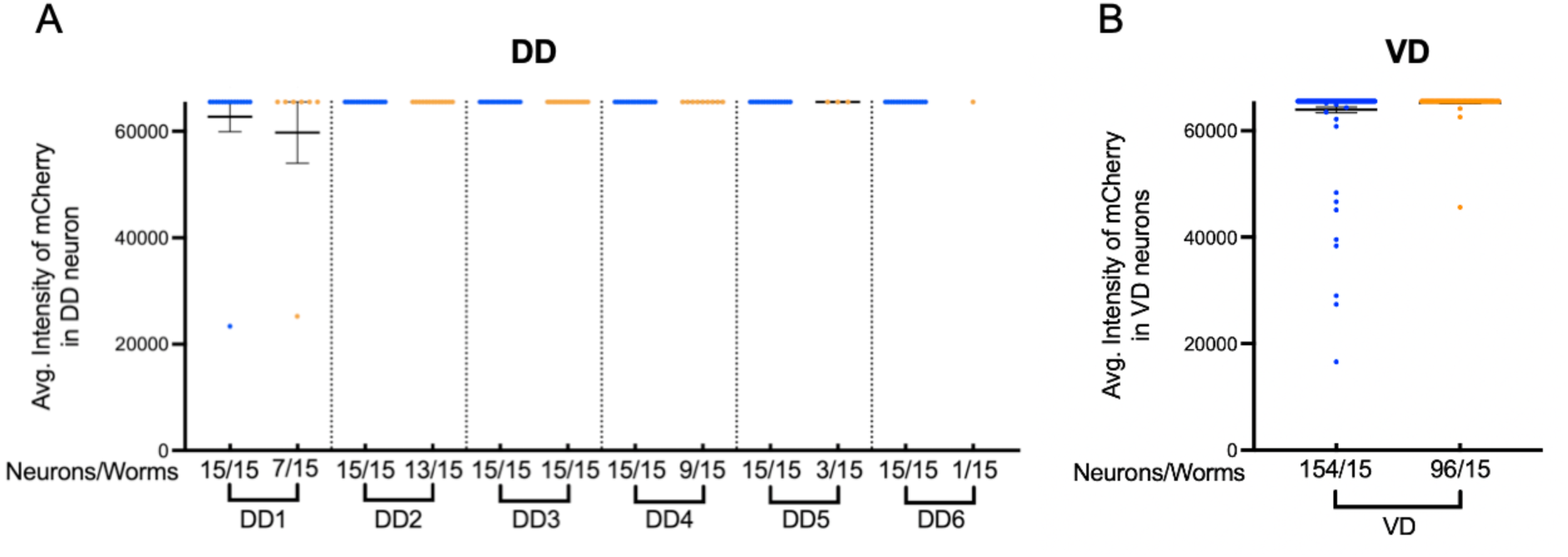
Effect of *smn-1* silencing on VD and DD Neurons Reporter expression. **(A-B)** Comparative analysis of *ttr-39p*::*mCherry* reporter intensity in each DD neuron (A) and VD neuron (B) at L4 larval stage in the control (blue dots) and in *smn-1*–silenced strains (orange dots).

**Figure S2.**
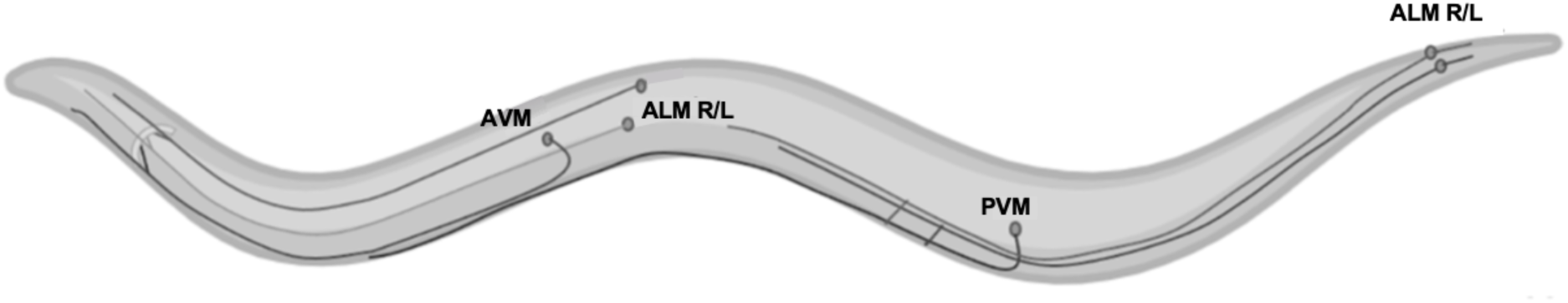
Anatomy of ALM and PLM neurite process. ALM (ALML, ALMR) neurons are located in the midbody and extend a long anterior process toward the pharynx, with a branch to the nerve ring. PLM (PLML, PLMR) neurons are located in the lumbar ganglion and extend a short posterior process and a long anterior process that ends near the ALM. AVM and PVM are positioned in the ventral body and extend a single long process along the ventral nerve cord.

### Supplementary Videos

**Supplementary Video S1.**
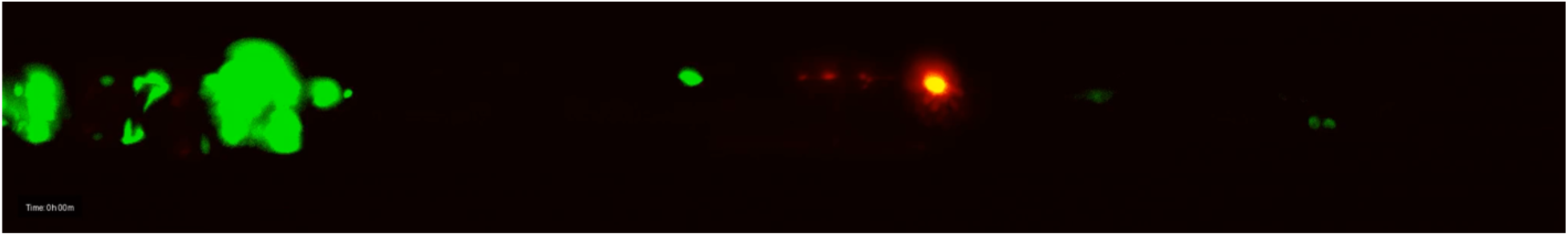
Long-term time-lapse imaging of D-type motor neurons in *smn-1*– silenced worms to monitor the birth of VD neurons (red neurons). During the L1-to-L2 transition, 4 out of 13 VD neurons in the complete set appeared. In this video, the DD3 neuron is visible expressing GFP alone, with no co-expression of mCherry. Time-lapse images were captured at 15-minute intervals.

**Supplementary Video S2:**
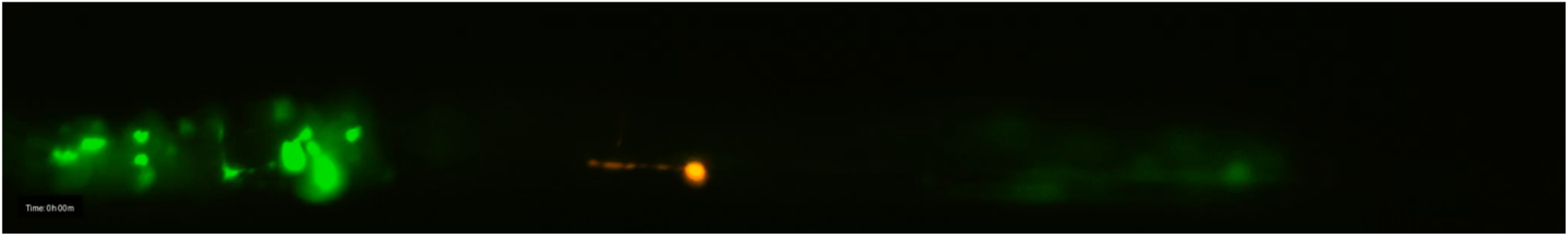
Long-term time-lapse imaging of D-type motor neurons in *smn-1*– silenced worms to monitor the birth of VD neurons (red neurons). During the L1-to-L2 transition, 9 out of 13 VD neurons in the complete set appeared. Time-lapse images were captured at 15-minute intervals.

**Supplementary Video S3:**
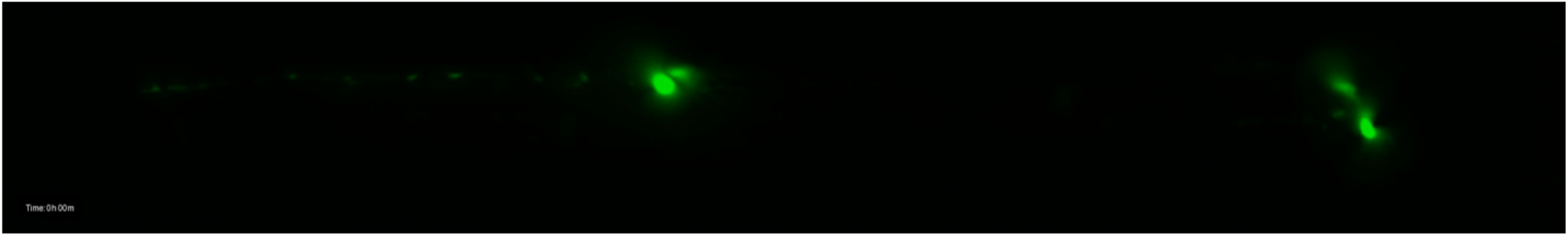
Long-term time-lapse imaging of PLMs in *smn-1*–silenced worms to monitor the survival of TRNs during larval stage. At the beginning of the video, both PLM neurons are visible (Two neurons in the right end of video). From ∼8 hours onward, one PLM neuron disappears, leaving only a single PLM neuron visible. Time-lapse images were captured at 15-minute intervals.

## Bibliography

Abati E, Citterio G, Bresolin N, Comi GP, Corti S. 2020. Glial cells involvement in spinal muscular atrophy: Could SMA be a neuroinflammatory disease? Neurobiology of Disease 140: 104870.

Anagnostou E, Miller SP, Guiot M-C, Karpati G, Simard L, Dilenge M-E, Shevell MI. 2005. Type I spinal muscular atrophy can mimic sensory-motor axonal neuropathy. J Child Neurol 20: 147– 150.

Bäumer D, Lee S, Nicholson G, Davies JL, Parkinson NJ, Murray LM, Gillingwater TH, Ansorge O, Davies KE, Talbot K. 2009. Alternative splicing events are a late feature of pathology in a mouse model of spinal muscular atrophy. PLoS Genet 5: e1000773.

Berger S, Spiri S, deMello A, Hajnal A. 2021. Microfluidic-based imaging of complete Caenorhabditis elegans larval development. Development (Cambridge, England) 148. https://www.ncbi.nlm.nih.gov/pmc/articles/PMC8327290/ (Accessed October 4, 2024).

Berger S, Spiri S, deMello A, Hajnal A, Berger S, Spiri S, deMello A, Hajnal A. 2025. High-Resolution C. elegans Imaging Across All Larval Stages. Journal of Visualized Experiments (JoVE) e68172.

Brenner S. 1974. The Genetics of CAENORHABDITIS ELEGANS. Genetics 77: 71–94.

Briese M, Esmaeili B, Fraboulet S, Burt EC, Christodoulou S, Towers PR, Davies KE, Sattelle DB. 2009. Deletion of smn-1, the Caenorhabditis elegans ortholog of the spinal muscular atrophy gene, results in locomotor dysfunction and reduced lifespan. Hum Mol Genet 18: 97–104.

Butchbach MER. 2016. Copy Number Variations in the Survival Motor Neuron Genes: Implications for Spinal Muscular Atrophy and Other Neurodegenerative Diseases. Front Mol Biosci 3: 7.

Chalfie M, Sulston J, White J, Southgate E, Thomson J, Brenner S. 1985. The neural circuit for touch sensitivity in Caenorhabditis elegans. J Neurosci 5: 956–964.

Chuang M, Goncharov A, Wang S, Oegema K, Jin Y, Chisholm AD. 2014. The microtubule minus end binding protein Patronin/PTRN-1 is required for axon regeneration in C. elegans. Cell Rep 9: 874–883.

Coer̈s C, Woolf AL. 1959. The innervation of muscle : a biopsy study. C.C. Thomas https://cir.nii.ac.jp/crid/1130282273373936640 (Accessed October 1, 2024).

Corti S, Nizzardo M, Simone C, Falcone M, Nardini M, Ronchi D, Donadoni C, Salani S, Riboldi G, Magri F, et al. 2012. Genetic Correction of Human Induced Pluripotent Stem Cells from Patients with Spinal Muscular Atrophy. Sci Transl Med 4: 165ra162.

Crawford TO, Pardo CA. 1996. The Neurobiology of Childhood Spinal Muscular Atrophy. Neurobiology of Disease 3: 97–110.

Doyle JJ, Vrancx C, Maios C, Labarre A, Patten SA, Parker JA. 2020. Modulating the endoplasmic reticulum stress response attenuates neurodegeneration in a Caenorhabditiselegans model of spinal muscular atrophy. Dis Model Mech 13: dmm041350.

Esposito G, Di Schiavi E, Bergamasco C, Bazzicalupo P. 2007. Efficient and cell specific knock-down of gene function in targeted C. elegans neurons. Gene 395: 170–176.

Faravelli I, Rinchetti P, Tambalo M, Simutin I, Mapelli L, Mancinelli S, Miotto M, Rizzuti M, D’Angelo A, Cordiglieri C, et al. 2025. Targeted antisense oligonucleotide treatment rescues developmental alterations in spinal muscular atrophy organoids. Nat Commun. https://www.nature.com/articles/s41467-025-67725-1 (Accessed January 16, 2026).

Gallotta I, Mazzarella N, Donato A, Esposito A, Chaplin JC, Castro S, Zampi G, Battaglia GS, Hilliard MA, Bazzicalupo P, et al. 2016. Neuron-specific knock-down of SMN1 causes neuron degeneration and death through an apoptotic mechanism. Hum Mol Genet 25: 2564–2577.

Hall DH, Russell RL. 1991. The posterior nervous system of the nematode Caenorhabditis elegans: serial reconstruction of identified neurons and complete pattern of synaptic interactions. J Neurosci 11: 1–22.

Hobert O. 2002. PCR fusion-based approach to create reporter gene constructs for expression analysis in transgenic C. elegans. Biotechniques 32: 728–730.

Howell K, White JG, Hobert O. 2015. Spatiotemporal control of a novel synaptic organizer molecule. Nature 523: 83–87.

Jablonka S, Karle K, Sandner B, Andreassi C, von Au K, Sendtner M. 2006. Distinct and overlapping alterations in motor and sensory neurons in a mouse model of spinal muscular atrophy. Hum Mol Genet 15: 511–518.

Jospin M, Qi YB, Stawicki TM, Boulin T, Schuske KR, Horvitz HR, Bessereau J-L, Jorgensen EM, Jin Y. 2009. A Neuronal Acetylcholine Receptor Regulates the Balance of Muscle Excitation and Inhibition in Caenorhabditis elegans. PLOS Biology 7: e1000265.

Kamath RS, Fraser AG, Dong Y, Poulin G, Durbin R, Gotta M, Kanapin A, Le Bot N, Moreno S, Sohrmann M, et al. 2003. Systematic functional analysis of the Caenorhabditis elegans genome using RNAi. Nature 421: 231–237.

Kirszenblat L, Neumann B, Coakley S, Hilliard MA. 2013. A dominant mutation in mec-7/β-tubulin affects axon development and regeneration in Caenorhabditis elegans neurons. Mol Biol Cell 24: 285–296.

Kong L, Valdivia DO, Simon CM, Hassinan CW, Delestrée N, Ramos DM, Park JH, Pilato CM, Xu X, Crowder M, et al. 2021. Impaired prenatal motor axon development necessitates early therapeutic intervention in severe SMA. Sci Transl Med 13: eabb6871.

Lotti F, Imlach WL, Saieva L, Beck ES, Hao LT, Li DK, Jiao W, Mentis GZ, Beattie CE, McCabe BD, et al. 2012. An SMN-dependent U12 splicing event essential for motor circuit function. Cell 151: 440–454.

Mariol M-C. 2013. A Rapid Protocol for Integrating Extrachromosomal Arrays With High Transmission Rate into the C. elegans Genome.

Martínez-Hernández R, Bernal S, Also-Rallo E, Alías L, Barceló MJ, Hereu M, Esquerda JE, Tizzano EF. 2013. Synaptic defects in type I spinal muscular atrophy in human development. J Pathol 229: 49–61.

McGovern VL, Gavrilina TO, Beattie CE, Burghes AHM. 2008. Embryonic motor axon development in the severe SMA mouse. Hum Mol Genet 17: 2900–2909.

Mello CC, Kramer JM, Stinchcomb D, Ambros V. 1991. Efficient gene transfer in C.elegans: extrachromosomal maintenance and integration of transforming sequences. EMBO J 10: 3959– 3970.

Mercuri E. 2021. Spinal muscular atrophy: from rags to riches. Neuromuscular Disorders 31: 998–1003.

Miguel-Aliaga I, Culetto E, Walker DS, Baylis HA, Sattelle DB, Davies KE. 1999. The Caenorhabditis elegans orthologue of the human gene responsible for spinal muscular atrophy is a maternal product critical for germline maturation and embryonic viability. Hum Mol Genet 8: 2133–2143.

Pellizzoni L, Yong J, Dreyfuss G. 2002. Essential role for the SMN complex in the specificity of snRNP assembly. Science 298: 1775–1779.

Ruggiu M, McGovern VL, Lotti F, Saieva L, Li DK, Kariya S, Monani UR, Burghes AHM, Pellizzoni L. 2012. A role for SMN exon 7 splicing in the selective vulnerability of motor neurons in spinal muscular atrophy. Mol Cell Biol 32: 126–138.

Simon CM, Dai Y, Van Alstyne M, Koutsioumpa C, Pagiazitis JG, Chalif JI, Wang X, Rabinowitz JE, Henderson CE, Pellizzoni L, et al. 2017. Converging mechanisms of p53 activation drive motor neuron degeneration in spinal muscular atrophy. Cell Rep 21: 3767–3780.

Singh RN, Howell MD, Ottesen EW, Singh NN. 2017. Diverse role of survival motor neuron protein. Biochim Biophys Acta Gene Regul Mech 1860: 299–315.

Spiri S, Berger S, Mereu L, DeMello A, Hajnal A. 2022. Reciprocal EGFR signaling in the anchor cell ensures precise inter-organ connection during Caenorhabditis elegans vulval morphogenesis. Development 149: dev199900.

Sulston JE, Schierenberg E, White JG, Thomson JN. 1983. The embryonic cell lineage of the nematode *Caenorhabditis elegans*. Developmental Biology 100: 64–119.

Tarrade A, Fassier C, Courageot S, Charvin D, Vitte J, Peris L, Thorel A, Mouisel E, Fonknechten N, Roblot N, et al. 2006. A mutation of spastin is responsible for swellings and impairment of transport in a region of axon characterized by changes in microtubule composition. Human Molecular Genetics 15: 3544–3558.

Thompson-Peer K, Bai J, Hu Z, Kaplan JM. 2012. HBL-1 patterns synaptic remodeling in C. elegans. Neuron 73: 453–465.

Toms N, Cooper J, Patchen B, Aamodt E. 2001. High copy arrays containing a sequence upstream of mec-3 alter cell migration and axonal morphology in C. elegans. BMC Dev Biol 1: 2.

Wen H-L, Lin Y-T, Ting C-H, Lin-Chao S, Li H, Hsieh-Li HM. 2010. Stathmin, a microtubule-destabilizing protein, is dysregulated in spinal muscular atrophy. Hum Mol Genet 19: 1766– 1778.

Werdnig G. 1891. Zwei frühinfantile hereditäre Fälle von progressiver Muskelatrophie unter dem Bilde der Dystrophie, aber anf neurotischer Grundlage. Archiv f Psychiatrie 22: 437–480.

White JG, Southgate E, Thomson JN, Brenner S. 1986. The structure of the nervous system of the nematode Caenorhabditis elegans. Philos Trans R Soc Lond B Biol Sci 314: 1–340.

Yeo CJJ, Darras BT. 2021. Yeo and Darras: Extraneuronal Phenotypes of Spinal Muscular Atrophy. Ann Neurol 89: 24–26.

Zhen M, Samuel ADT. 2015. C. elegans locomotion: small circuits, complex functions. Curr Opin Neurobiol 33: 117–126.

Zheng C, Jin FQ, Trippe BL, Wu J, Chalfie M. 2018. Inhibition of cell fate repressors secures the differentiation of the touch receptor neurons of Caenorhabditis elegans. Development 145: dev168096.

